# Bronco T (*Shirisadi kasaya*), a polyherbal formulation ameliorates LPS induced septicemia in rats

**DOI:** 10.1101/2021.10.23.465588

**Authors:** Priyanka Mishra, Ratna Pandey, Suyash Tripathi, Sushil K Dubey, Yamini B Tripathi

## Abstract

Septicemia is a life-threatening state, leading to multi-organ failure, ARDS and death. So, efforts are being made to identify novel therapies. Here, Bronco T (BT), a polyherbal formulation developed in 1984 for treating asthma, has been repurposed against septicemia induced ALI. The LPS (3mg/kg BW) was injected intraperitoneally before 24 hours, of surgery to assess the cardiorespiratory parameters, blood PaO_2_/FiO_2_, pulmonary water content and histological changes in the lungs. The pentoxifylline (PTX) (25 mg/kg b.w.) was used as the positive control. The PTX was given one hour before LPS and BT was given 3 hours (orally in different doses of 3, 1.5 and 0.75 gm/kg BW) to maintain the Cmax of the drug. The LPS treated group showed significant bradypnea, bradycardia and low heart rate frequency as observed, through elongated peaks (RR) and (MAP) respectively and finally death after 95 minutes of LPS injection. The PTX and BT (3gm/kg) pretreatment significantly prevented these changes (dose-dependent in the BT group). The survival was maintained up to 190 min after LPS. The Pentoxifylline showed a better response (75%) than Bronco T (72%). In both the treatments, a significant decrease in pulmonary water content and minimal neutrophil infiltration and intact alveoli-capillary membrane was seen in the transverse section (T.S) of the lungs. **Conclusion**: Significant improvement was noted in survival time, lesser tissue damage and better lung physiology by treating with Bronco T in LPS induced septicemia.

## Introduction

Septicemia is initiated by a host-mediated inflammatory response during infection and in uncontrolled conditions, resulting in ARDS and death even after ventilator support, in intensive care units (ICUs)(1-3). Sepsis development has two phases: the 1^st^ phase involves an overwhelming burst of pro-inflammatory cytokines, targeted to inhibit the growth of pathogens, but its long time persistence causes pathological damage. (4). In the 2nd phase, the host’s immune response is partially deregulated, resulting in an imbalance between pro-inflammatory and anti-inflammatory cytokine responses that ultimately leads to life-threatening tissue damage, multiple organ failure, and vascular dysfunction and death. Sepsis is frequently associated with ARDS characterized by severe respiratory failure with reduced pulmonary functioning and increased oxidative stress(5). As sepsis progresses the heart starts beating rapidly and breathing becomes rapid as a compensatory mechanism in the initial stage but followed by a drop in blood pressure (6), resulting from reduced oxygen supply to vital organs like the brain, heart, lungs and kidney leading to multiple organ failure(7). Despite the use of efficient anti-inflammatory therapies and multi-organ function support, the mortality rate is still significantly high. The patients with septicemias, hospitalized in ICU on ventilator support also have a poor prognosis, as per the APACHE III score. The oxygenation index is compromised in patients with sepsis and ARDS due to hypoxemia and respiratory distress(8). Histological examination in these patients showed hyaline membranes, and mixed infiltration of inflammatory cells in the interstitium, alveoli, and perivascular areas (9).

Gram-negative bacterial infection is one of the major causes of septicemia mediated ALI which is attributed to Lipopolysaccharide (LPS) mediated activation of TLR4 receptor and downstream signalling pathway for high synthesis of inflammatory cytokines like IL6 and TNF alpha.

The Pentoxifylline (1-(5-oxohexyl)-3,7-dimethylxanthine) has been used as the positive control therapy. It is a competitive nonselective phosphodiesterase inhibitor. It increases cellular cAMP level, activates PKA, inhibits TNF and leukotriene synthesis and down-regulates inflammation and innate immunity cascade.

The Bronco T is a polyherbal decoction developed by late Prof. S.N.Tripathi for the management of asthma and it is in clinical use since 1984. It consists of obtained from five plants including Sirisha (Albizia lebbeck), Kantkari (Solanum virginianum), Vasaka (Justicia adhatoda), Madhuyasthi (Glycyrrhiza glabra), and Tejpatra (Cinnamomum tamala), having therapeutic claims against the upper respiratory tract disorders(10)(11). However, no investigation has been done, either with BT or its constituent plants for the management of ALI/ARDS. Here, for the 1^st^ time, we have tested this new therapeutic claim of BT on an experimental model of sepsis-induced ALI, to analyze its effect on cardio-respiratory, biochemical and histological parameters.

## Material and Method

### 2.1 Drugs preparation and administration

Urethane and Lipopolysachharides (*Escherichia coli* 0111:B4, Lot # 019M4009V) were obtained by Sigma Aldrich Inc, St Louis, USA. Pentoxifylline being sold as Trentrol 400 tablet (400mg per tablet, Batch No.INA0007, Sanofi), and Bronco-T (70gm, Batch No-161, Surya Pharmaceuticals) were purchased from the local market.

### 2.2 Methodology

#### 2.2.1 Experimental Protocol

The Animal Ethical Committee of IMS-BHU approved the current study (letter No. Dean/2021/IAEC/2550). The inbred adult *albino* rats of Charles-Foster strain weighing (175- 225 g) were purchased from our central animal house and acclimatized for 7 days in ambient conditions of temperature, and a day-light cycle of 12 hrs with food and water ad libitum. Urethane 1.5 g/kg body weight i.p.) was given to anaesthetize the animals. An additional bolus dose of urethane (0.1-0.15 g/kg i.p.) was injected if required. The Trachea, jugular vein and carotid artery were cannulated to maintain the respiration and for drug administration, and record the blood pressure, respectively. The animals were stabilized for 30 min after dissection and cannulation. The animals were divided into six groups (n = 6).

| S.No | Groups | Vehicle | Route |
| --- | --- | --- | --- |
| 1 | Group I (Experimental Control) | Normal Saline (75µl) | Peroral |
| 2 | Group II (Disease control) | LPS (3mg/kg) 24 hours | Intraperitoneal |
| 3 | Group III (Positive control) | Pentoxifylline (25mg/kg) i.p. 1 hour before LPS injection. | Intraperitoneal |
| 4 | Group IVa (BT-Treatment) | (3gm/kg) orally 3 hours before LPS injection and after 6 hours. | Peroral |
| 5 | Group IVb (BT-Treatment) | (1.5 gm/kg) orally 3 hours before LPS injection and after 6 hours. | Peroral |
| 6 | Group IVc (BT-Treatment) | (0.75 gm/kg) orally 3 hours before LPS injection and after 6 hours. | Peroral |

**Note: The dose and duration of treatment were made based on the biological half-life of the active constituents**.

At the end of each experiment, the lungs were excised, and one lung was kept for the estimation of pulmonary water content and the other one was used for histological examination. Blood from the right internal carotid artery was collected for blood gas analysis.

#### 2.2.2 Treatment

Pentoxifylline was pre-administered at the dose of 25mg/kg b.w. i.p to the rats one hour before LPS administration (3mg/kg, 24 hours). The BT decoction was orally given three hours before LPS and after six hours to maintain a steady Cmax of BT. The selected doses were 3,1.5 and 0.75 gm/kg of rat body weight. (11). This dose was calculated, based on a human dose i.e one teaspoon thrice a day for an adult (60Kg), which corresponds to approximately 3.196 gm, which is equivalent to 150mg/kg of human body weight. The rat dose was 10 times higher than the human dose because rats have a metabolic rate, about ten times higher than humans (12)(13). The selected ranges of 3 doses were 0.75gm, 1.5gm and 3gm/kg of BW of rats for experimental study.

### 2.3. Protocol for parameter estimation

#### 2.3.1 Determination of survival time

It was estimated for the observation period of 190 min as the LPS treated group died around 95 min after 24 hours, therefore the rationale for twice the survival time of the LPS group was chosen. Mortality was characterized by flat line response in cardio-respiratory responses (RR, MAP, and HR) as observed from the Lab Chart recorder (AD Instrument).

#### 2.3.2 Determination of Respiratory Rate

It was recorded by securing the skin over the xiphisternum with help of a thread and connecting it to the chart recorder (AD Instrument) through a force-displacement transducer.

#### 2.3.3 Determination of Mean Arterial Pressure (MAP)

It was estimated from blood pressure, firstly the pressure transducer was calibrated and the carotid artery was connected to it through a three-way stop clock. Blood pressure was recorded by connecting the pressure transducer to a chart recorder (AD Instrument) through a bridge amplifier. MAP was determined by computing the data obtained.

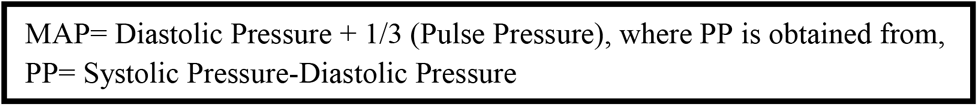

#### 2.3.3 Determination of Heart Rate (HR)

It was manually calculated from R-R interval by recording the electrocardiograph by connecting the needle electrode to the chart recorder (AD Instrument) through bio amplifier using standard limb lead II configuration in the rats.

#### 2.3.4 Determination of PaO_2_/FiO_2_ratio

PaO_2_/FiO_2_ ratio was estimated by ABG analyzer (Roche OMNI gas analyzer) by collecting 100µl blood sample from right internal carotid artery using a heparinized syringe.

#### 2.3.5 Determination of pulmonary water content

It was determined by the physical method as described earlier(14). At the end of each experiment, the lungs were excised. One lung was preserved in formal saline for histological examination and the other was weighed and dried in an electric oven (at 90°C for 48 h) to a constant weight. The difference between wet weight and dry weight was calculated to determine the water content.

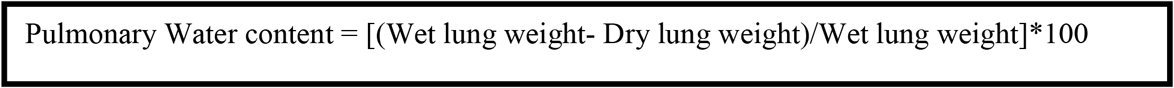

#### 2.3.6 Histology of Lungs

Surface morphology was observed and photographed, at the end of the experiment. The lung tissue was washed with ice-cold phosphate buffer saline and preserved in a 10% formalin solution. Further, it was subjected to standard histological protocol and 5-micron lung tissue sections were stained with hematoxylin (H) and eosin (E) for microscopic examination (10X, 40X magnification)

## Result

### A) Effect of Bronco T and PTX in LPS treated rats on physiological parameters

Bronco T (3gm/kg) significantly improved the physiological parameters in LPS treated rats. On observation rats in Group, I moved freely after 24 hours and did not show any sign of respiratory distress. However, LPS treated rats were comparatively less active and showed shortness of breath. BT pretreated rats were significantly in better condition with 100 per cent survival.

#### 1) Survival Time

All the animals in Group-I survived throughout the experimental period (190 min). In LPS treated group-II, the survival time was reduced to **92 min +/-0.3**. Further, in Pentoxifylline (25mg/kg) treated group-III, survival time was significantly higher than the LPS treated group, but lower than NC. In the BT treated group 3 gm/kg vs 25 mg IV and V, dose-dependent enhancement in survival time was recorded, however, in group VI (0.75mg/kg) the survival time was 110 min **+/-0.3**. (**Fig 2A-F)**.

**Fig 2A-F:**
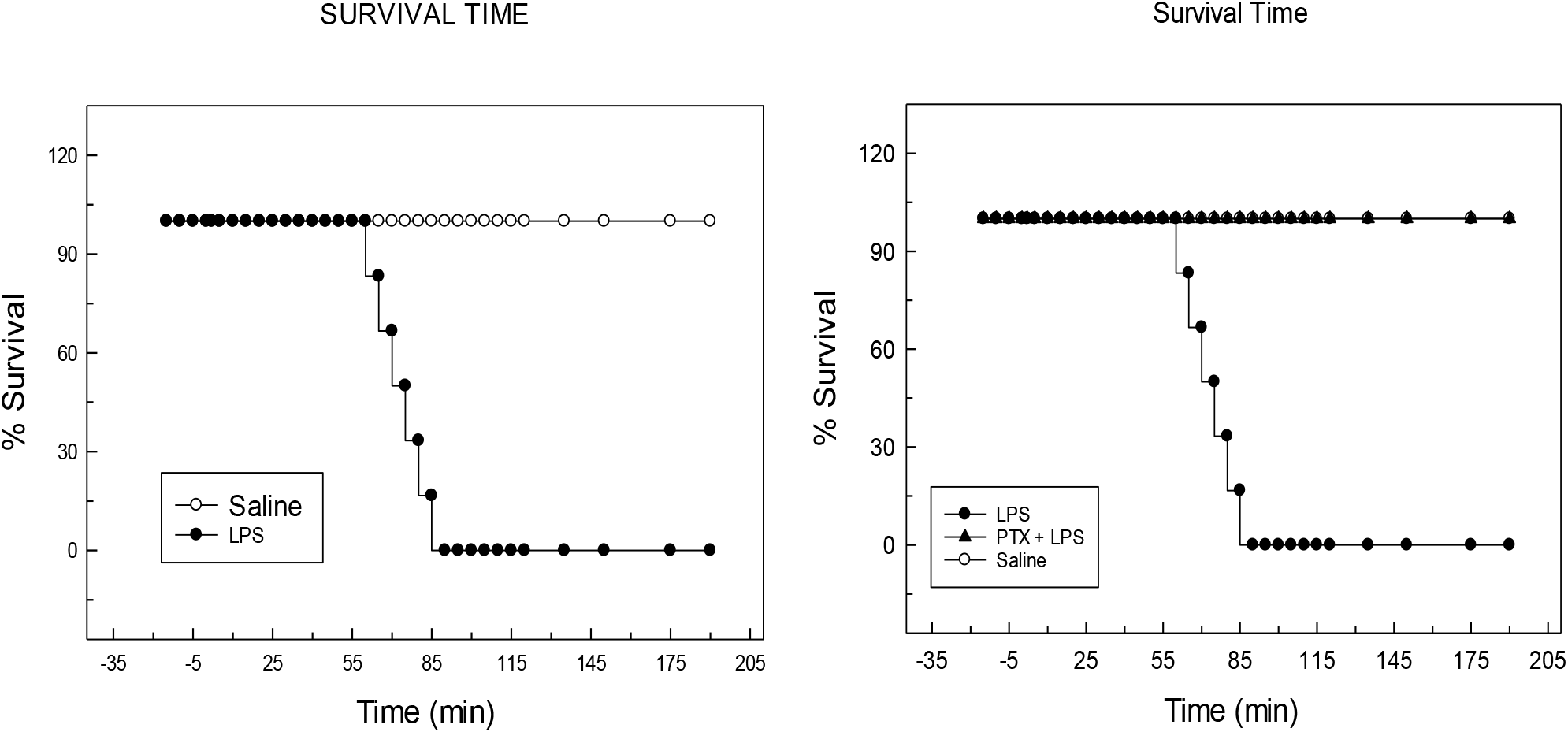
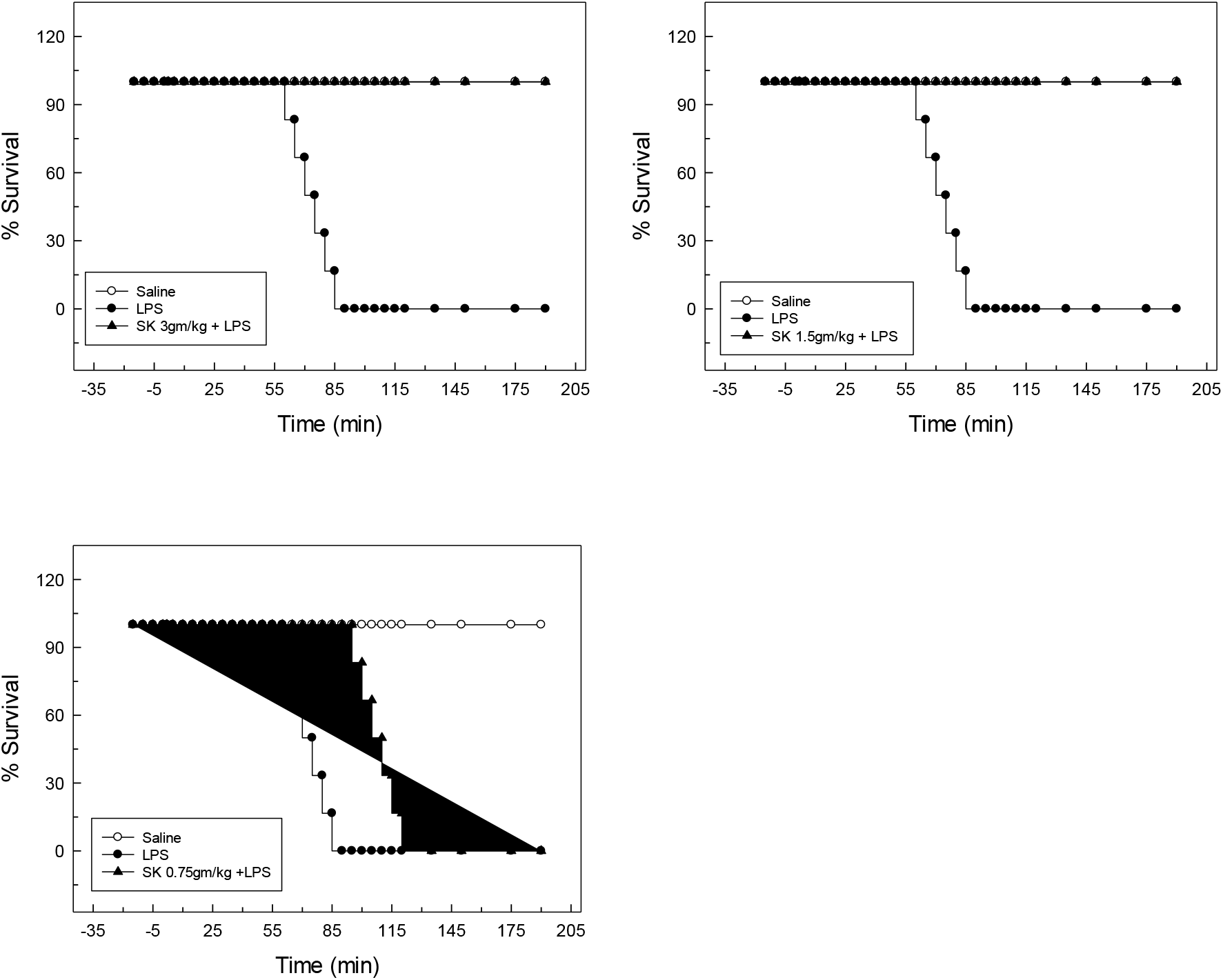
Survival time comparison in each group as obtained from Kaplein Meyers plot.2A denotes comparison between normal saline and LPS.2B denotes comparison between LPS and PTX.2C-E denotes comparison between LPS and BT (3,1.5 and 0.75gm/kg BW).

#### 2) Cardiorespiratory parameters

In group I, after 24 hours and surgical stabilization, no significant change was observed in the RR, MAP and HR. However, in group II there was a significant reduction in RR, followed by death after 90 minutes. Interestingly, in group III, the RR was reduced in the beginning but later on maintained throughout the observation period. In the BT pretreatment group, RR was maintained to data of group I (normal saline-treated rats) (Fig 3A). A similar observation was noted on HR (Fig 3B) and MAP (Fig 3C) parameters.

**Fig3A:**
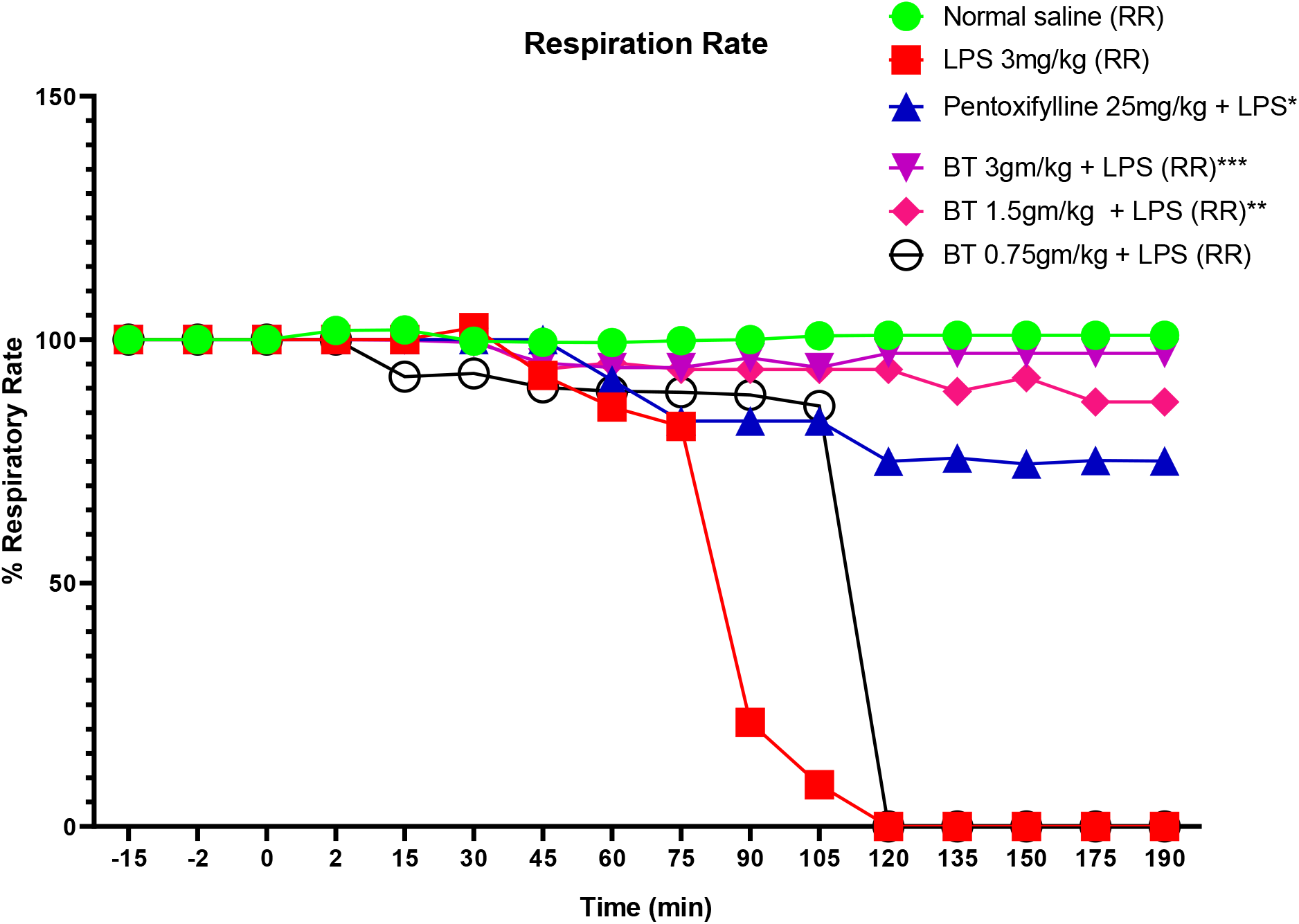
Respiration Rate in each group. LPS treated group suffered bradypnea from starting and represented flat line response after 95 minutes (Red blocks).BT treated group (3gm/kg) showed significant improvement in comparison to PTX treated group. Data represented as percentage initial Mean +/-SD where ***, p<0.001,** p<0.01, * p<0.05 in comparison to LPS 3mg/kg treated group. Note: (-) 15, indicates the time taken for stabilization of animal post-surgical interventions

**Fig3B:**
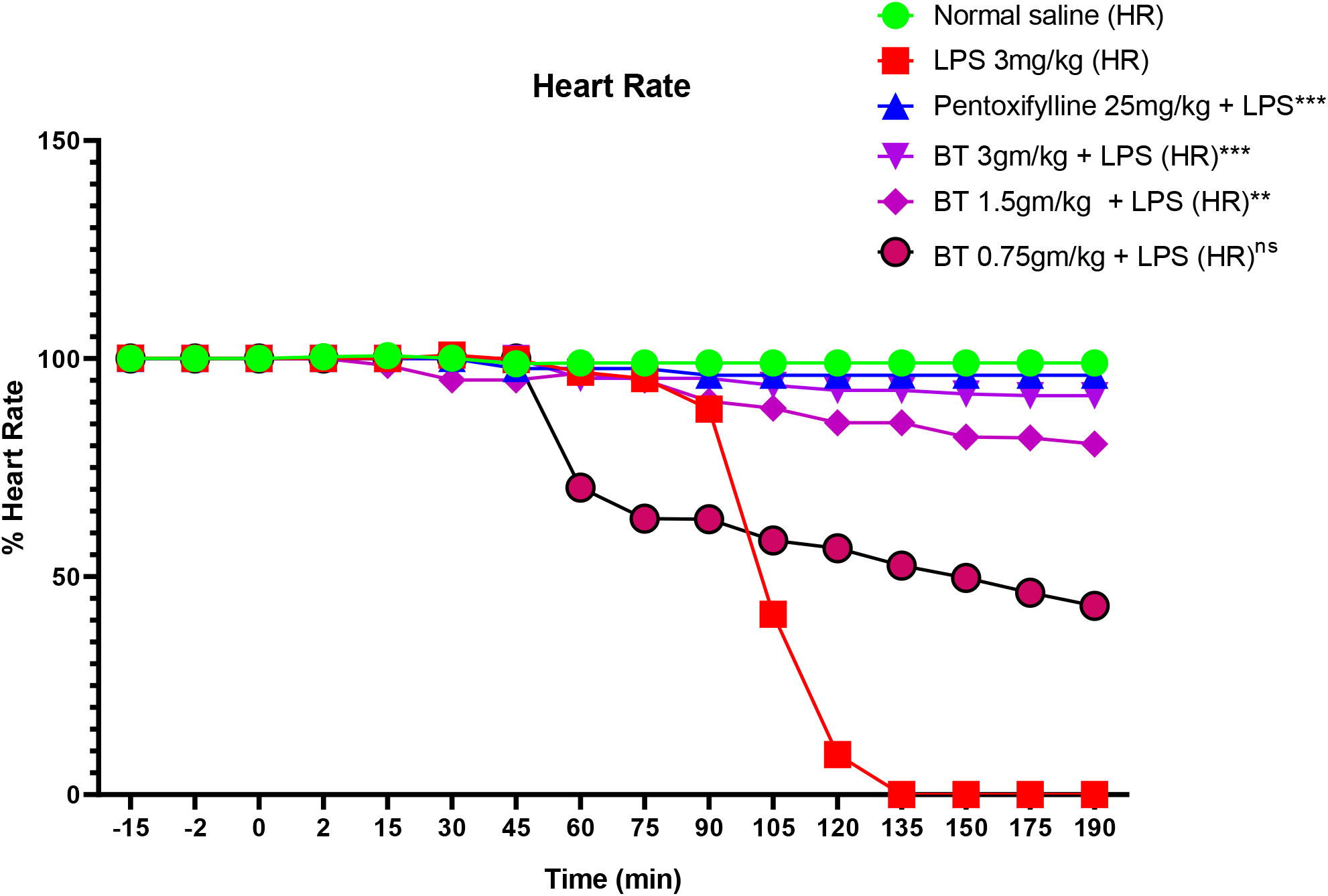
Heart Rate in each group. LPS treated group suffered bradycardia from starting and represented flat line response after 120 minutes (Red blocks).BT treated group (3gm/kg) showed significant improvement in comparison to PTX treated group. Data represented as percentage initial Mean +/-SD where ***,p<0.001,** p<0.01, * p<0.05 in comparison to LPS 3mg/kg treated group.

**Fig3C:**
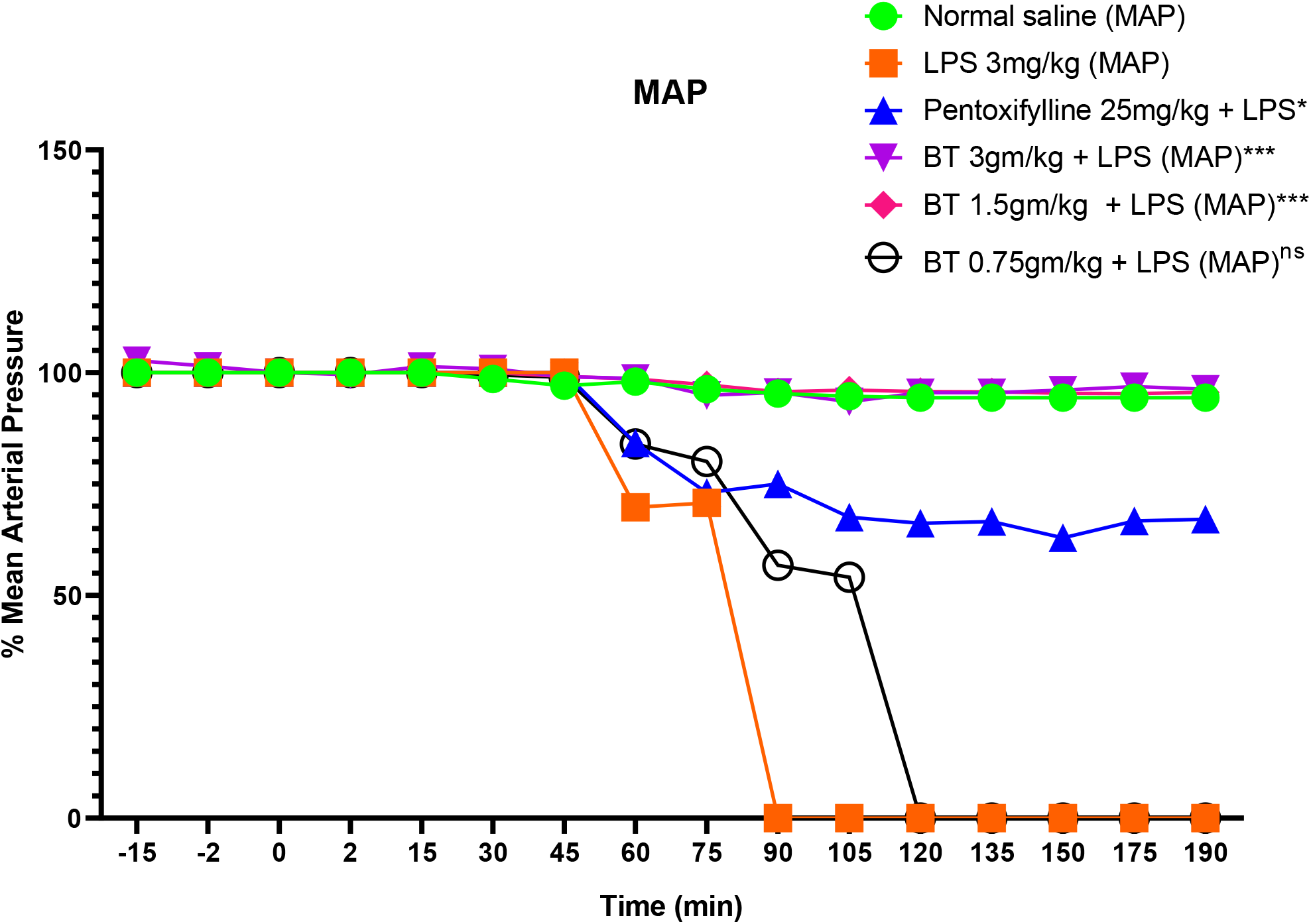
Mean arterial pressure in each group. LPS treated group represented flat line response after 75 minutes (Orange blocks).BT treated group (3gm/kg) showed significant improvement in comparison to PTX treated group. Data represented as percentage initial Mean +/-SD where ***,p<0.001,** p<0.01, * p<0.05 in comparison to LPS 3mg/kg treated group.

**Fig 4:**
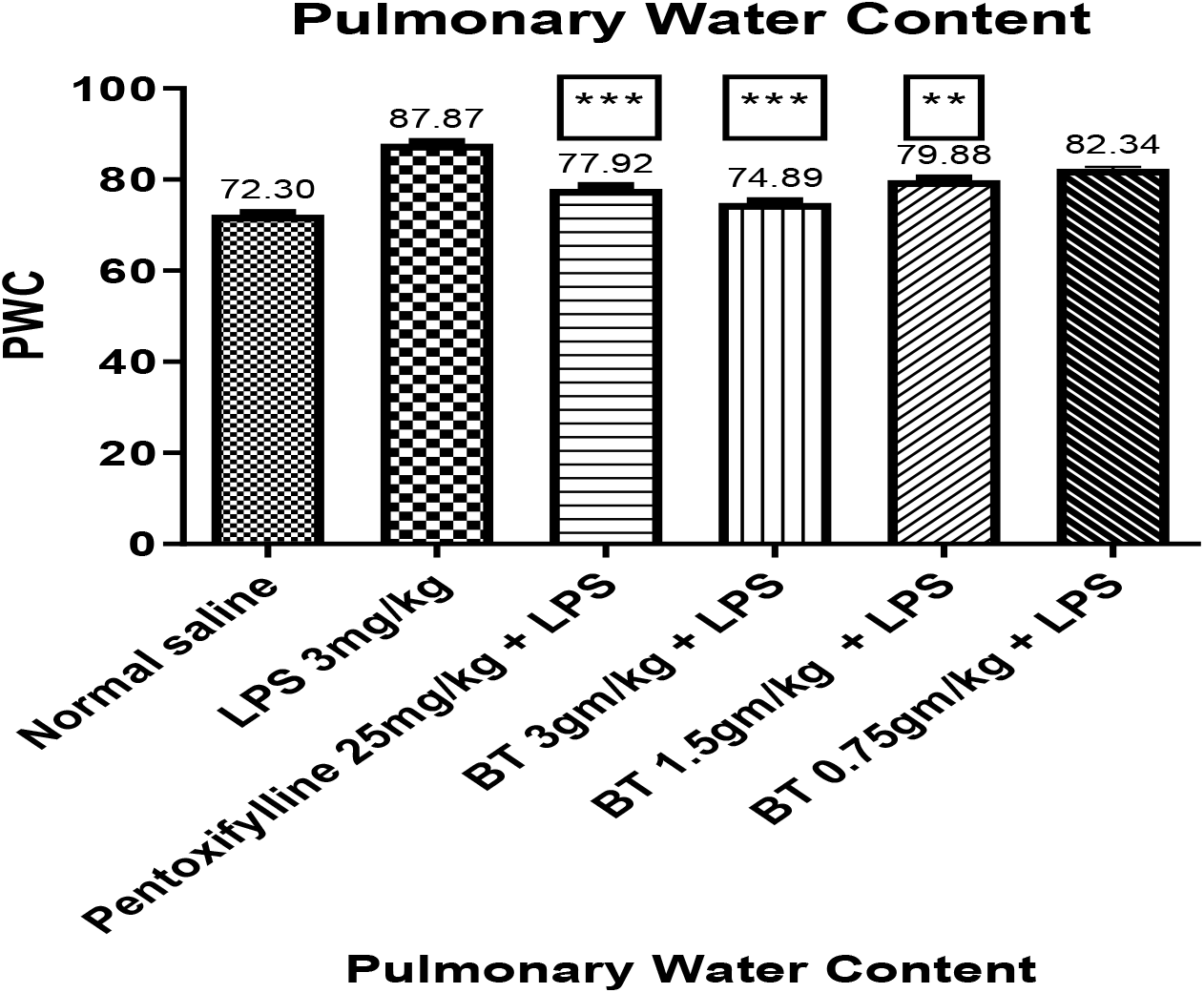
Effect on lung wet to dry ratio in each group. Here BT (3 gm/kg) showed significant improvement in comparison to PTX treated groups. Data represented as Mean +/-SD where ***,p<0.001,** p<0.01, * p<0.05 in comparison to LPS 3mg/kg treated group following one-way ANOVA Turkey test.

#### 3) PaO_2_:FiO_2_ (Oxygenation Index)

The PaO_2_:FiO_2_ represent the oxygenation capacity of the lungs, which indirectly determines the mortality and directly determine the ARDS incidence. The hypoxemic condition will lead to respiratory distress and multi-organ failure if not treated. As shown in **Table 2**, LPS treated group have PaO_2_:FiO_2_ less than 300 signifying the acute lung injury condition. This was increased in both PTX and BT (3gm/kg) treated groups suggesting their preventive role against acute lung injury.

**Table2:**
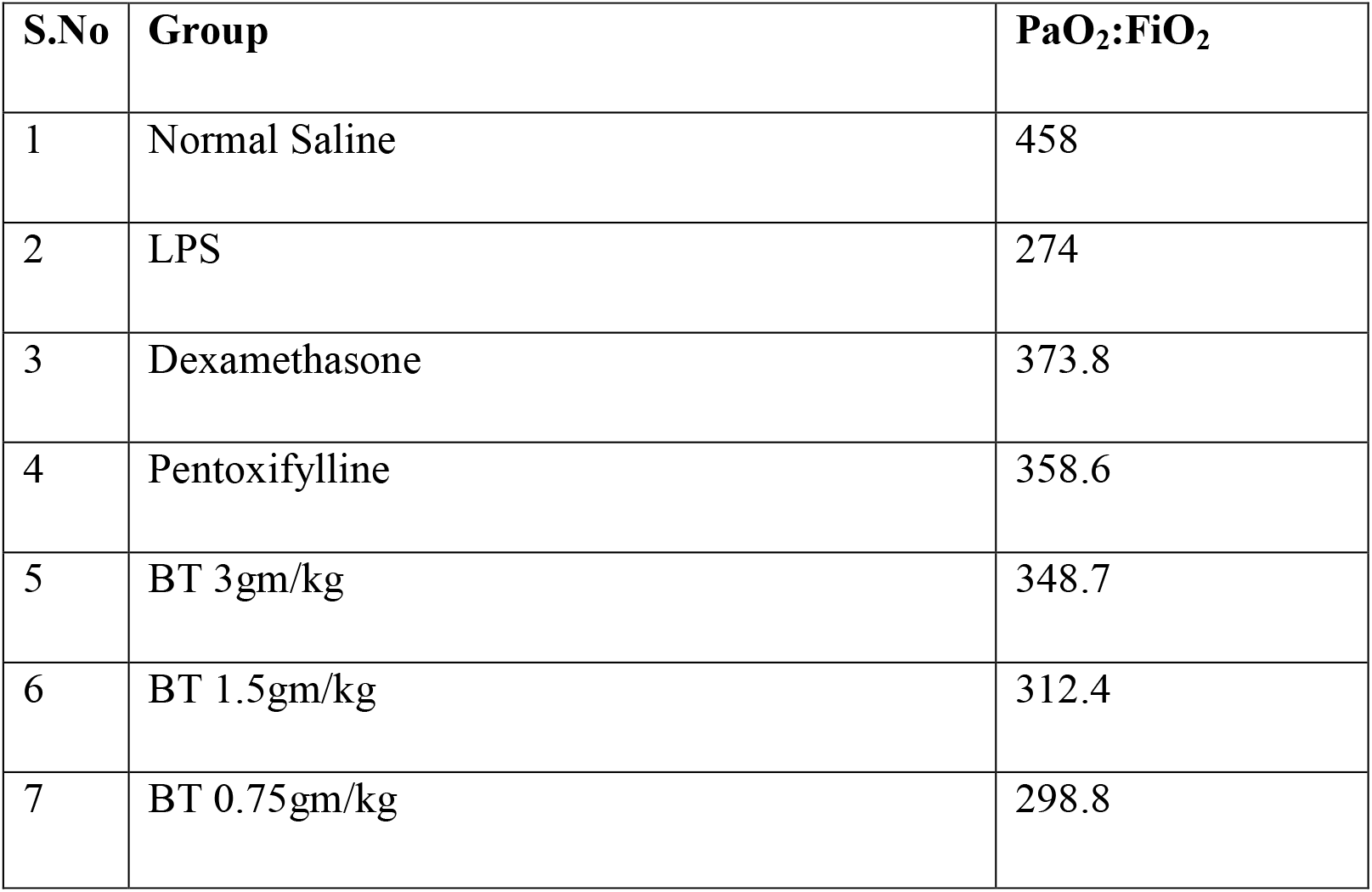
Oxygenation Index (PaO_2_:FiO_2_) in each group.

### B) Effect of Bronco T and PTX in LPS treated rats on pulmonary parameters

#### 1) Lung Wet to Dry ratio (Pulmonary water content)

PWC indicates the level of oedema in the pulmonary region, the higher the water content higher will be fluid accumulation and reduced lung compliance. The wet to dry ratio for all the treated groups were determined after 24 h following the LPS challenge. A significant increase in lung W/D weight ratio was observed for the LPS group as compared to normal saline. However, BT pre-treatment (3 gm/kg) significantly reduced lung W/D weight ratios (n = 6, p < 0.001) as compared to the LPS group. Similarly, the PTX treated group also markedly reduced the wet to dry weight ratio compared to the LPS as shown in the figure below.

#### 2) Surface Morphology

The surface morphology analysis revealed **(Fig 5)** in group I, the tissue was dry and without hyperemia, however, in group II, the surface appeared to be dark red to black with a hardened and congested appearance a typical characteristic of ALI. The morphological presentation was better and there was significantly less bleeding in the PTX (group III) and BT (group IV) groups.

**Fig 5:**
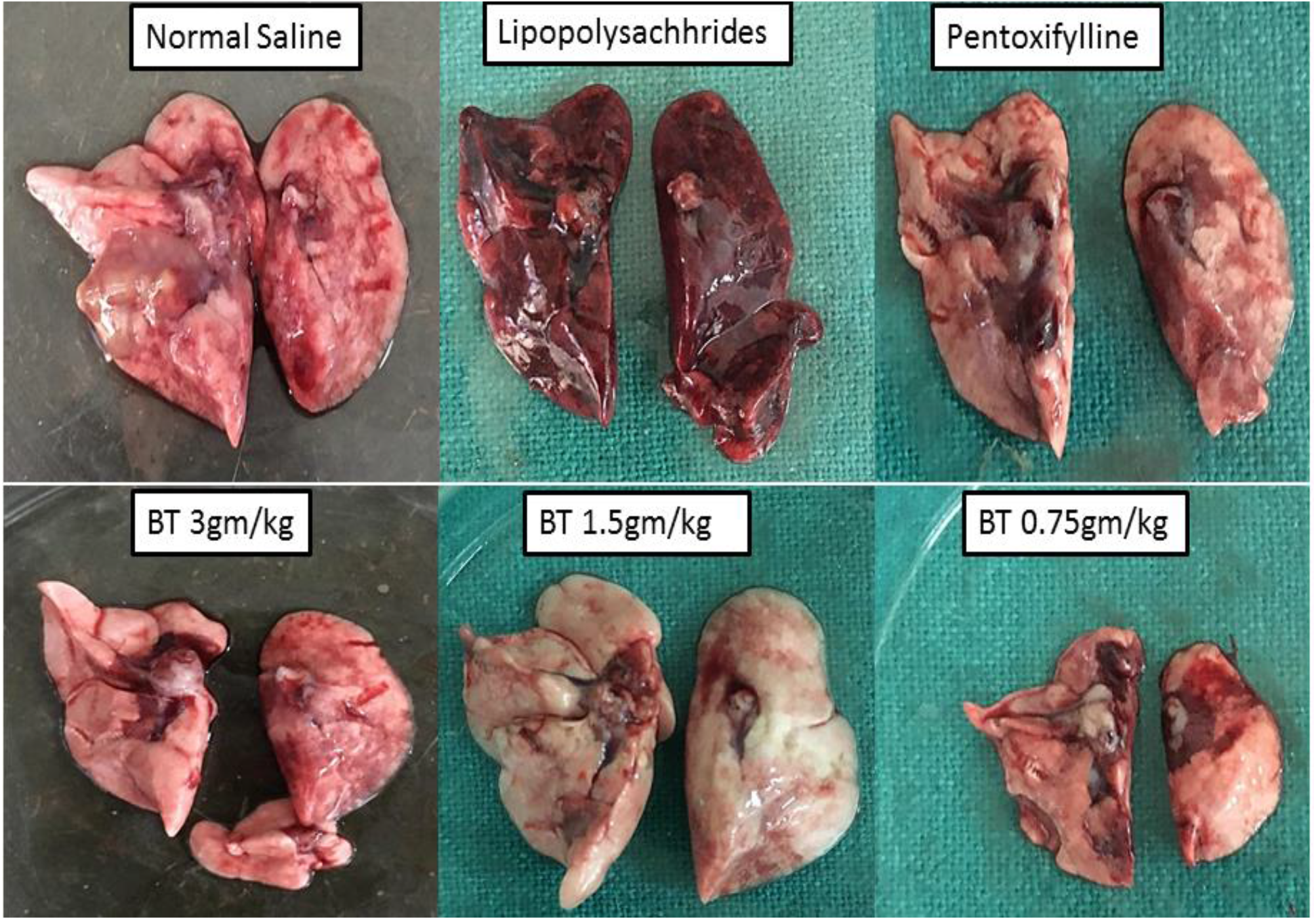
Surface morphology of lungs of each group.

#### 3) Histological examination

To reveal the pathological changes H& E staining was used. As shown in **Fig 6 (10x) and 7(40x)** LPS administration was associated with significantly more fluid accumulation and inflammatory cells in the alveolar region. There is thickened alveolo-capillary membrane and hemorrhagic manifestation in this group. In contrast, BT and PTX treated groups showed varying degrees of protection against LPS.BT (3gm/kg BW) treated group have fewer cell infiltration due to intact alveolo-capillary membrane. Their histological appearance is closer to the group I.In the figure A denotes group I(NS),B denotes group II (LPS),C denotes group III (PTX 25mg/kg BW),D denotes group IV (BT 3gm/kg BW),E denotes group IV (BT 1.5gm/kg) and F denotes group IV (BT 0.75 gm/kg BW) respectively.

**Fig 6:**
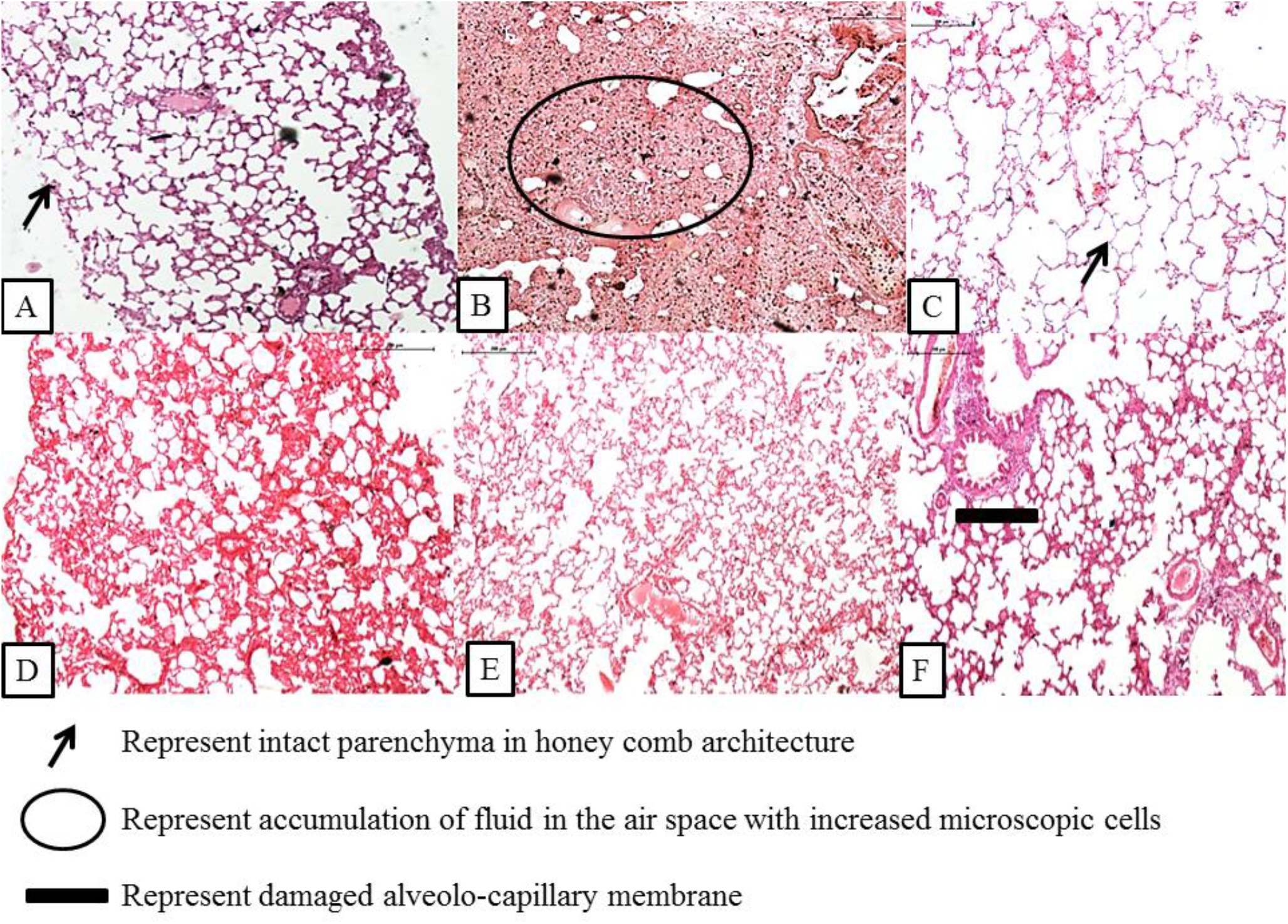
Histological section (H&E staining) of lungs in each group at 10X magnification reveals the following information-. 1) In Group I(A) normal honeycomb presentation of parenchymal cells could be observed which in Group II(B) is flooded with protein-rich fluid and there is clear sloughing of the alveolar region. 2) In Group III (C) PTX treatment prevented the sloughing of the alveolar region but honeycomb presentation is altered 3) In Group IV (D, E, F) clear airspace are visible with near to normal histoarchitecture which prevents the stiffening of lungs

**Fig 7:**
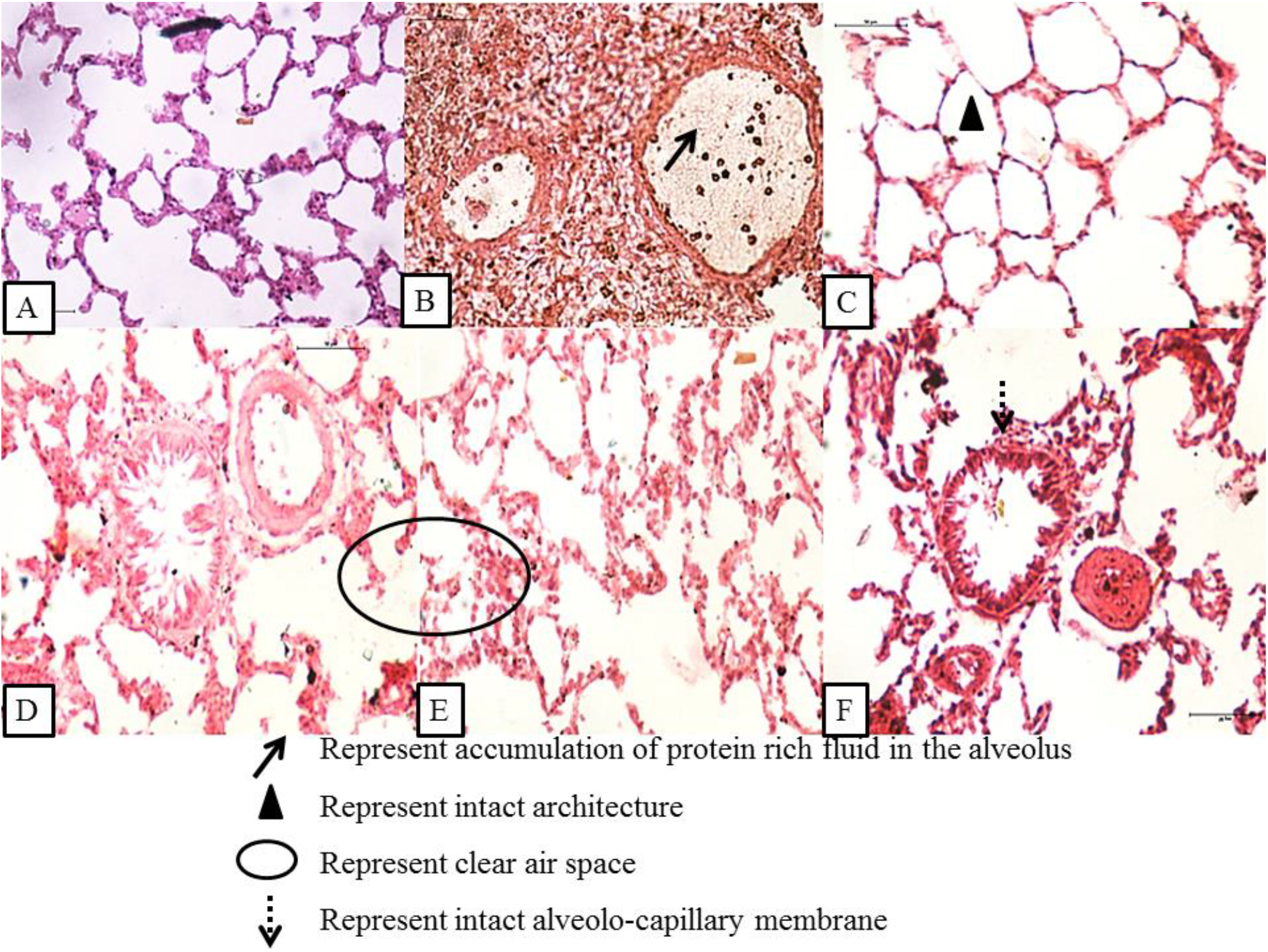
Histological section of lungs in each group at 40X magnification reveals the following information-. 1) In Group II (B) the alveolo-capillary membrane is breached with the intrusion of neutrophils and fluid accumulation in this region 2) In Group III (C) the intact structure of parenchymal tissue could be observed. 3) In Group IV (D, E, F) clear airspace and intact endothelial layer could be observed. There is no hemorrhagic manifestation with less number of intruding cells in the alveolar region.

## Discussion

Septicemia is the leading cause of mortality due to its respiratory distress and multi-organ failure characteristics. Since pentoxifylline are currently the only dependable pharmacological intervention in sepsis(15), so we have used it as a positive control for comparison of the efficacy of BT on sepsis-related physiological, pulmonary and histological changes. Since the heart-lung interaction is difficult to evaluate during mechanical ventilation in sepsis-induced ARDS patients (16), so here we have estimated these in rats. In the present work, we have demonstrated the protective role of Bronco T in LPS induced sepsis in rats on the parameters described above. In septicemia, initially, the respiratory rate is increased as protection of animal in acute condition, and then gradually reduces which is deleterious if not managed timely. This reduced respiration-rate leads to lowering of oxygen in the pulmonary region and cause severe hypoxemia. In this study, we observed that after 24 hours of LPS administration, there was a decrease in respiration rate per min with 2-3 peaks per five seconds. Pentoxifylline treatment reverses this altered effect by inhibiting pro-inflammatory cytokines including TNF alpha and leukotriene synthesis(17). Similarly, BT 3gm/kg BW treatment significantly reverse this effect with near to normal peak presentation suggesting that the herbal decoction was able to save the lungs from the deleterious effect of LPS.

Mean Arterial Pressure (MAP) is the main hemodynamic variable indicating the driving pressure for organ perfusion. In the patient with septicemia, all blood vessels dilate due to the release of nitric oxide, causing blood pressure to drop. Infection coupled with lack of blood flow to vital organs leads to septic shock and organ failure(20)(21). PTX was able to maintain MAP>65mmHg due to inhibition of nitric oxide(18) and improving hemodynamic capacity(19). Similar observations were found with BT 3gm/kg treatment indicating the same mechanism of action which need to be further explored.

In severe sepsis, initially, the heart beats rapidly, followed by bradycardia where the heart is unable to pump blood to vital organs irrespective of any interventions(22). Here in our study, a similar presentation was found in LPS treated group where there was persistent hypotension and bradycardia during recording after 24 hours. PTX and BT had maintained the heart rate to the normal range of 96.7% and 91.5% respectively. These suggest that BT had a cardio protective role against LPS in a similar manner to PTX.

The ratio of the partial pressure of arterial oxygen to fraction of inspired oxygen (P/F) determines the degree of severity of sepsis. It quantifies severe hypoxemic respiratory failures(23). Hypoxemia presents various complications including an elevated level of inflammatory cytokines, reactive oxygen species, neutrophils and reduced lung compliance. LPS treatment for 24 hours lowered the P/F ratio to severe grade. This was rectified in BT and PTX treated group implicating the role of BT against hypoxemia and its detrimental effect.

Septicemia produces symptoms of pneumonia due to increased water content. Increased pulmonary water content leads to poor lung compliance and indirectly plays a key role in the failure of the respiratory system. This happens due to altered membrane permeability and inactivation of Na/K+ ATPase pump (24). The lung wet to dry ratio determines this condition objectively. LPS treatment increased lung wet to dry ratio, therefore, stiffens the lung and leading to poor exchange of gases of the lungs. BT treated groups significantly lowered the lung wet to dry ratio suggesting it maintain the integrity of the alveoli-capillary membrane and activates epithelial Na/K+ ATPase pump in LPS treated rats. However, more study requires pinpointing the exact mechanism behind this.

This physiological change has been further supported by histological studies where we found there is a peculiar presentation of hyaline membrane formation, with an increased number of inflammatory cells in the interstitium and alveolar area(9). This is a typical illustration of ARDS due to injured Type I and II pneumocytes which leads to insufficient surfactant production. This was visible in LPS treated rats where the lungs were hardened with hemorrhagic manifestation. There was an increased fluid accumulation and number of infiltrating cells in the lungs. This was rectified in BT and PTX treated rats significantly. On histological analysis, there was intact endothelial and epithelial barrier in BT treated group along with clear air space and less number of infiltrating cells. This indicates the lung-protective role of BT in LPS treated rats in comparison to PTX treatment.

## Conclusion

From analyzing the above-stated parameters, it can be safely suggested that Bronco T (BT) significantly improved survival time, cardiorespiratory, physiological, pulmonary and histological parameters in the LPS induced animal model of septicemia. Therefore BT can be used as add on therapy with all treatments, recommended in the guidelines of conventional therapies, even in patients, admitted on mechanical support. Further study is required to elucidate the molecular mechanism of this preparation.

## Acknowledgement

Priyanka Mishra is thankful to IMS-BHU for providing resources to carry out this work.

